# IGF2BP2 links m6A-modified HMGA1 mRNA stability to mitochondrial metabolism and cell proliferation

**DOI:** 10.64898/2026.04.24.720611

**Authors:** Zhao-Ming Cui, Jiang-Tao Lu, Yuan-Ye Xu, Ying-Ying Feng, Yang Guo, Yan-Ping Gao, Wen Wang, Le-Le Qiu, Xiao-Yin Wang, Zi-Chun Hua, Tian-Yun Wang, Yan-Long Jia

## Abstract

Chinese hamster ovary (CHO) cells serve as the primary host for industrial therapeutic protein production, yet enhancing their productivity remains a significant challenge. Epigenetic regulation, particularly RNA N6-methyladenosine modification, offers a promising strategy. Here, we show that the m6A reader protein insulin-like growth factor 2 mRNA-binding protein 2 (IGF2BP2) positively regulates recombinant protein yield in CHO cells. IGF2BP2 expression was elevated in high-producing clones, and its stable overexpression promoted cell proliferation, increased the S-phase cell proportion, and boosted titers and specific productivity of recombinant proteins—adalimumab, vitronectin, and donanemab—by 2.0-, 1.6-, 2.6-fold and 1.8-, 1.4-, 2.1- fold, respectively. Mechanistically, IGF2BP2 recognized m6A sites on HMGA1 mRNA, enhancing its stability and expression. Integrated analyses of oxidative stress, mitochondrial function, and metabolomics, along with inhibitor validation, revealed that IGF2BP2 also strengthens antioxidant defense, promotes mitochondrial ATP production and utilization, and reshapes cellular redox and metabolic homeostasis. These findings highlight IGF2BP2 as a critical regulator of recombinant protein expression and mitochondrial oxidative metabolism in CHO cells, illustrating how RNA methylation cooperates with mitochondrial function and proliferation to enhance protein production.

## Introduction

Chinese hamster ovary (CHO) cells have become the most widely used host system in the biopharmaceutical field, owing to their excellent genetic stability and capabilities for proper protein folding and post-translational modifications; they are extensively applied in the production of recombinant therapeutic proteins and monoclonal antibodies (Zhang et al, 2022; Baek et al, 2025; Geng et al, 2024). However, challenges such as unstable transgene expression, clonal variation, and environmental stresses during large-scale culture (e.g., nutrient depletion, metabolite accumulation) persistently limit production efficiency, where these issues contribute to increased production costs (Martínez et al, 2024; Zhang et al, 2025). epigenetic regulation presents new opportunities to construct high-performance CHO cell lines through targeted cellular modification. This approach enhances the efficiency and sustainability of biopharmaceutical manufacturing without altering the underlying DNA sequences (Wang et al, 2024).

N6-methyladenosine (m6A) modification, the most prevalent epigenetic modification in eukaryotic messenger RNA, has been demonstrated in recent years to play a pivotal role in regulating gene expression (Zhang et al, 2025). As a dynamic and reversible RNA modification, m6A participates extensively in biological processes, including cell differentiation and metabolic homeostasis. This occurs through a “write-erase-read” regulatory mechanism involving methyltransferases (e.g., METTL3 and METTL14), demethylases (e.g., FTO and ALKBH5), and recognition proteins (e.g., the YTHDF family and IGF2BPs) (Sun and Zhang, 2021; Yang et al, 2023; Zhang et al, 2025). Insulin-like growth factor 2 mRNA-binding protein 2 (IGF2BP2), a crucial m6A “reader” protein, specifically recognizes and binds to m6A sites to enhance target mRNA stability and promote cellular proliferation (Zhao et al, 2020; Zhang et al, 2024; Shi et al, 2023). Specifically, IGF2BP2 binds to the m6A-modified region of SLC1A5, a key gene in glutamine metabolism, approximately doubling its mRNA half-life. This ensures sustained expression of SLC1A5, thereby promoting abnormal proliferation in tumor cells (Pu et al, 2025). Furthermore, in acute myeloid leukemia, IGF2BP2 enhances mRNA stability by binding m6A-modified sites on genes such as MYC and GPT2, sustaining their robust expression to regulate cellular proliferation and metabolism (Weng et al, 2022). Similarly, this “m6A modification-IGF2BP2 binding-mRNA stability” regulatory pattern occurs in colorectal cancer, where the IGF2BP2 complex likely promotes HMGA1 expression by recognizing its m6A-modified sites, thereby enhancing mRNA stability and facilitating cell cycle progression (Hou et al, 2021).

The chromatin regulator and transcriptional co-activator high mobility group A1 (HMGA1) promotes proliferation through multiple mechanisms. As a co-activator, HMGA1 binds DNA and recruits factors such as STAT1 to activate proliferation-associated genes including DDAH1 and GATA2 (Hu et al, 2025; Li et al, 2022). Its own expression is regulated by upstream transcription factors such as TFAP2A (Zhao and Lan, 2023). In cardiomyocytes, HMGA1 activates proliferation-related gene programs by increasing chromatin accessibility and reducing repressive histone marks like H3K27me3 (Bouwman et al, 2025). HMGA1 also functions within the PI3K/AKT pathway: it enhances gene expression and supports cell survival in response to external stimuli (Wang et al, 2022), and upregulates transcription of FOXO1—a downstream target of AKT—thereby promoting cell-cycle progression and inhibiting apoptosis (Xie et al, 2021).

Given the established role of IGF2BP2 in cell proliferation, redox homeostasis, and metabolic reprogramming (Zeng et al, 2025; He et al, 2025), and the absence of studies on m6A-mediated regulation of HMGA1 in CHO cells, this work investigated whether modulating IGF2BP2 expression could enhance recombinant protein production. We demonstrate that IGF2BP2 stabilizes HMGA1 mRNA by binding to specific m6A sites, upregulating HMGA1 expression. Elevated HMGA1 activates the PI3K/AKT/mTOR pathway to drive proliferation, ultimately increasing antibody titers. Moreover, IGF2BP2 overexpression reprograms cellular metabolism, improving glucose and glutamine utilization. This metabolic shift redirects flux from lactate production to consumption, reducing inhibitory byproduct accumulation, improving redox and energy balance, and enhancing mitochondrial function. Our findings reveal an IGF2BP2-mediated epigenetic mechanism that promotes both cell growth and metabolic fitness via HMGA1 stabilization and downstream proliferative signaling, providing a novel strategy for constructing high-performance CHO production cell lines.

## Results

### 1. Differential expression of IGF2BP2 in heterogeneous recombinant CHO cells

Our prior study reported variations in m6A modifications and transcriptomes among monoclonal clones exhibiting CHO cell heterogeneity (Wang et al, 2025). Since the relationship between overall expression levels of m6A-reading proteins in CHO cell lines and recombinant protein yield remains unclear, we systematically assessed mRNA expression levels of all IGF2BP family members in the recombinant CHO cells with high-/low- expression of adalimumab (ADM^H^ versus ADM^L^). Results showed that within the IGF2BP family, only IGF2BP2 exhibited significant differences in mRNA expression (Figure 1A,B). Subsequent Western blot and immunofluorescence analyses further demonstrated thatIGF2BP2 expression levels positively correlated with recombinant antibody expression levels (Figure 1C-F). These findings suggest that IGF2BP2 function and regulation may directly influence recombinant protein expression in CHO cells.

**Figure 1.**
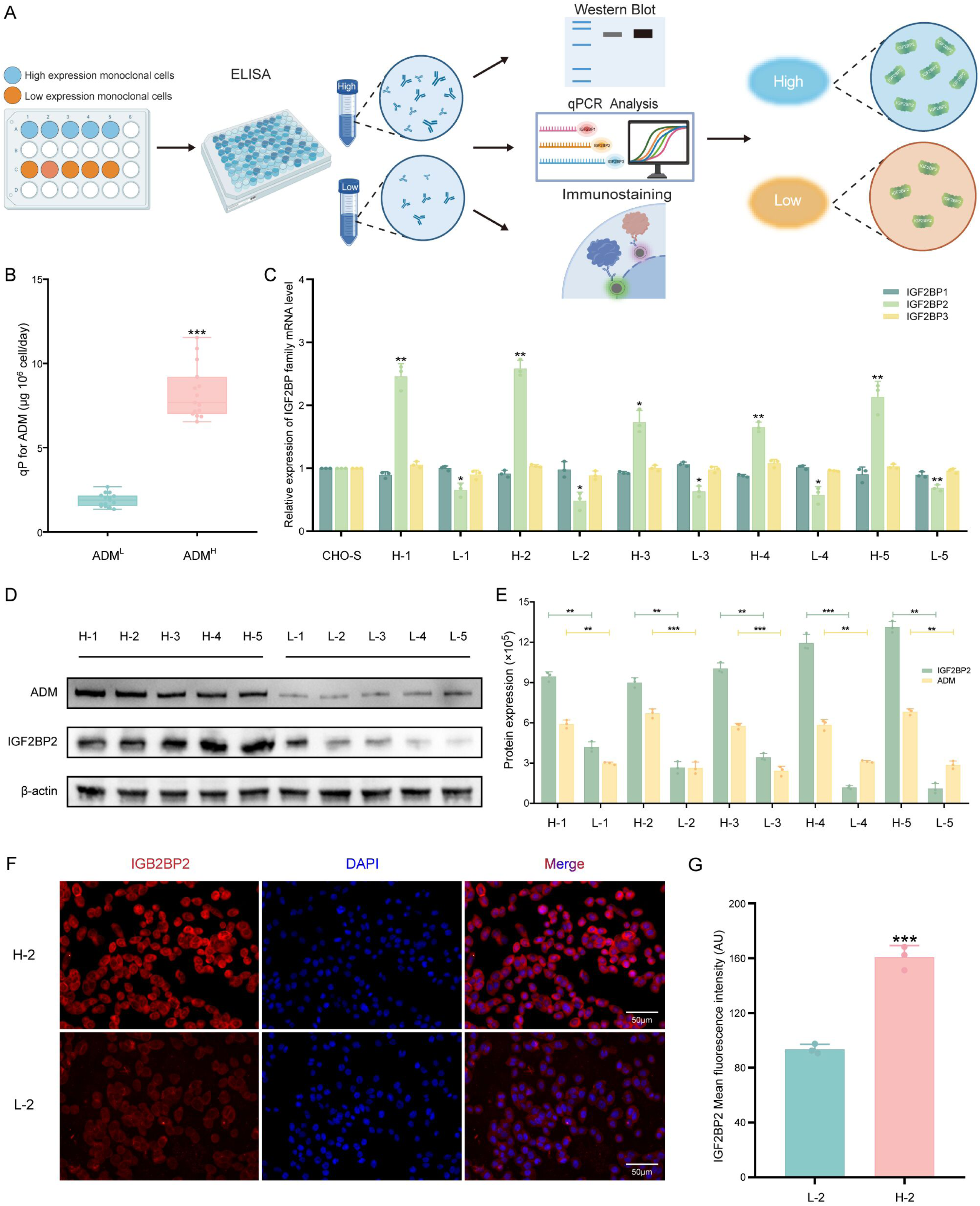
The expression levels of rADM demonstrated a significant positive correlation with IGF2BP2 expression. (A): Experimental workflow for screening IGF2BP2 as a key regulator of recombinant protein expression in CHO cells. (B): Relative protein expression of ADM quantified using ELISA. Relative mRNA expression levels of IGF2BP family genes were assessed via qPCR (C), while protein expression of IGF2BP2 and rADM was evaluated by Western blot (D, E). (F): Immunofluorescence images of CHO cells labeled with an anti-IGF2BP2 antibody (red) and counterstained using DAPI (blue). (G): Mean fluorescence intensity (AU) values of IGF2BP2. (*p < 0.05; **p < 0.01; ***p < 0.001)

### 2. Establishment of stable IGF2BP2-overexpressing CHO cell line and their effect on recombinant protein production

To establish a stable IGF2BP2-overexpressing cell line, normal CHO cells (CHO-S) transfected with the IGF2BP2-expression vector underwent a three-weekselection process under hygromycin pressure. qPCR and Western blot analyses confirmed significantly increased IGF2BP2 expression at the mRNA and protein levels, and immunofluorescence further verified the successful generation of the stable cell line OE IGF2BP2 (Figure 2A-C). Functional assays indicated that IGF2BP2 overexpression promoted cell proliferation, as evidenced by an increased proportion of cells in S phase, without significantly altering apoptosis levels (Figure 2D, F). To further investigate the effect of IGF2BP2 overexpression on proliferation, CCK-8 and colony formation assays were conducted. These results confirmed a marked enhancement in proliferation for the OE IGF2BP2 group compared to the control CHO cells (Figure 2E, G). Subsequently, three recombinant protein expression vectors (rADM, rVN, and rDONA) were transfected into the cells of both experimental and control groups for suspension culture, respectively. While cell viability remained comparable during the culture period, the OE IGF2BP2 group displayed a significantly higher viable cell density after the logarithmic growth phase (day 4) (Figure 2H, J, L). Finally, ELISA test results demonstrated significantly increasedyields of rADM (2.04±0.23-fold), rVN (1.62±0.18-fold), and rDONA (2.55±0.52-fold), with specific productivity (qP) elevated by 1.81±0.04-fold, 1.40±0.07-fold, and 2.10±0.05-fold, respectively, relative to the control group (Figure 2I, K, M).

**Figure 2.**
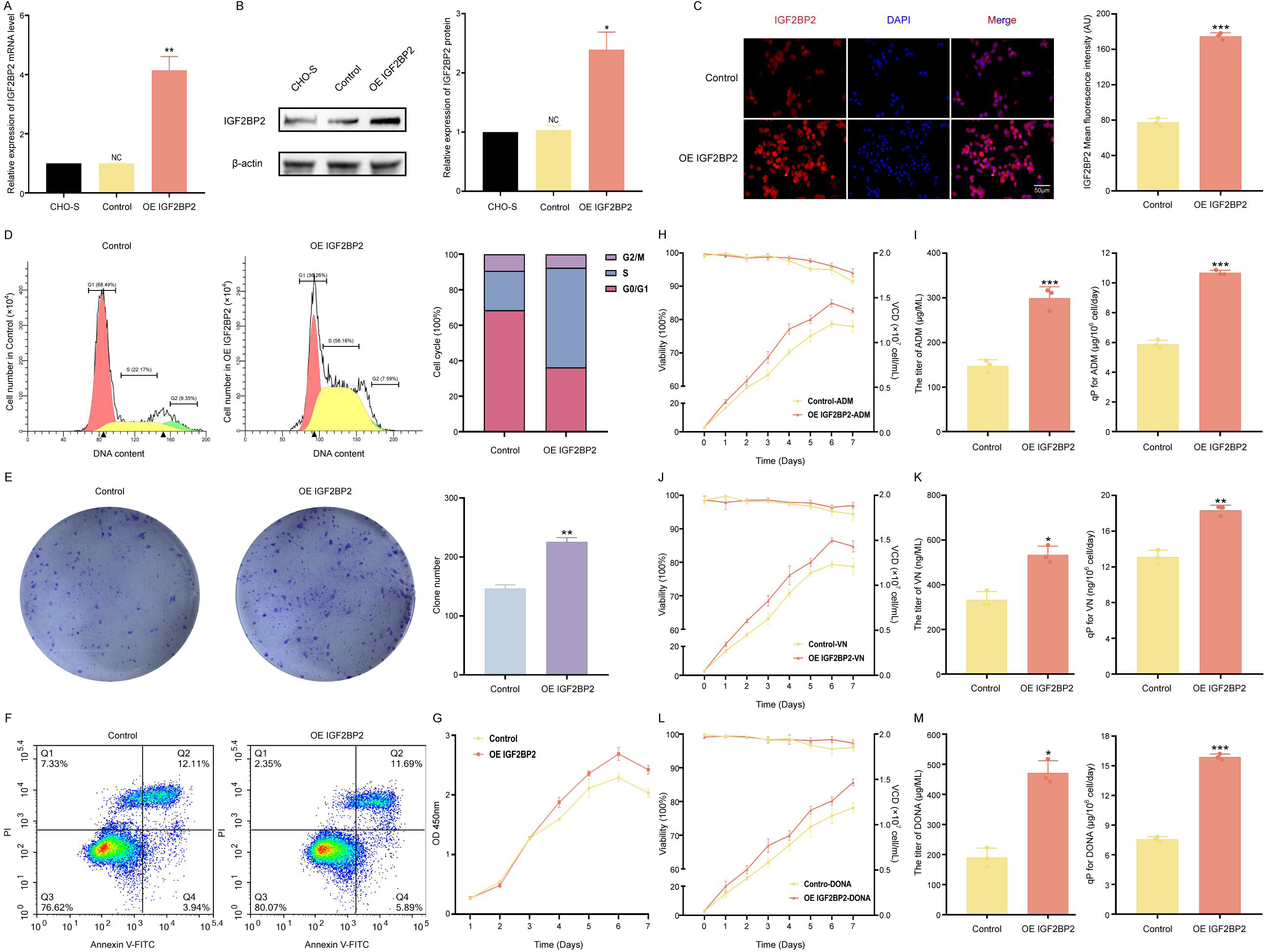
Expression of IGF2BP2, biological characterization of CHO cells transfected with an IGF2BP2 overexpression vector (OE IGF2BP2 group) compared with untransfected control cells (control group), and the subsequent effects on recombinant protein production. (A) IGF2BP2 mRNA expression was quantified using qPCR, and its protein levels were evaluated by Western blot analysis (B). (C) Immunofluorescence staining was performed on CHO cells using an anti-IGF2BP2 antibody (red), with nuclei counterstained with DAPI (blue); the average fluorescence intensity (in arbitrary units, AU) was also calculated. (D) Cell cycle distribution was analyzed. (E) Clonogenic potential was assessed via colony formation assay. (F) Apoptosis was measured to evaluate cell death. (G) Cell proliferation was determined using the CCK-8 assay. Viability and viable cell density (VCD) of CHO cells were examined following treatment with rADM (H), rVN (J), or rDONA (L) in both groups. The titer and quantitative production (qp) of rADM (I), rVN (K), and rDONA (M) were evaluated by ELISA in the two experimental groups. (*p < 0.05; **p < 0.01; **p < 0.001)

### 3. Rearrangement patterns and epigenetic modifications of differentially expressed genes (DEGs) in IGF2BP2-overexpressing CHO cells

To explore potential mechanisms enhancing recombinant protein expression after IGF2BP2 overexpression, we conducted RNA sequencing to assess global transcriptome changes and identify affected downstream signaling pathways. Volcano plots identified significantly DEGs (Figure 3A), which may function as key epigenetic regulators in the CHO cell recombinant protein production system. KEGG and GO analyses indicated that IGF2BP2 overexpression significantly enriched processes including activation of proliferation-related pathways (PI3K-AKT and mTOR) and positive regulation of cell proliferation (Figure 3B,C). Integrating the observed S-phase-upregulated cellular phenotype (Figure 2D), we cross-referenced DEGs from three related GO terms (Mitotic Cell Cycle, G1/S Transition of Mitotic Cell Cycle, Regulation of Cell Population Proliferation) and identified nine target genes, including HMGA1, Ppard, Stat6, and Ezh2 (Figure 3D). Subsequent GeneMANIA analysis of gene interaction networks revealed that HMGA1 may directly regulate IGF2BP2 (Figure 3E). Furthermore, heat maps, GSEA, and IGV analyses related to proliferation collectively corroborated that IGF2BP2 promotes cellular proliferation (Figure 3F-H).

**Figure 3.**
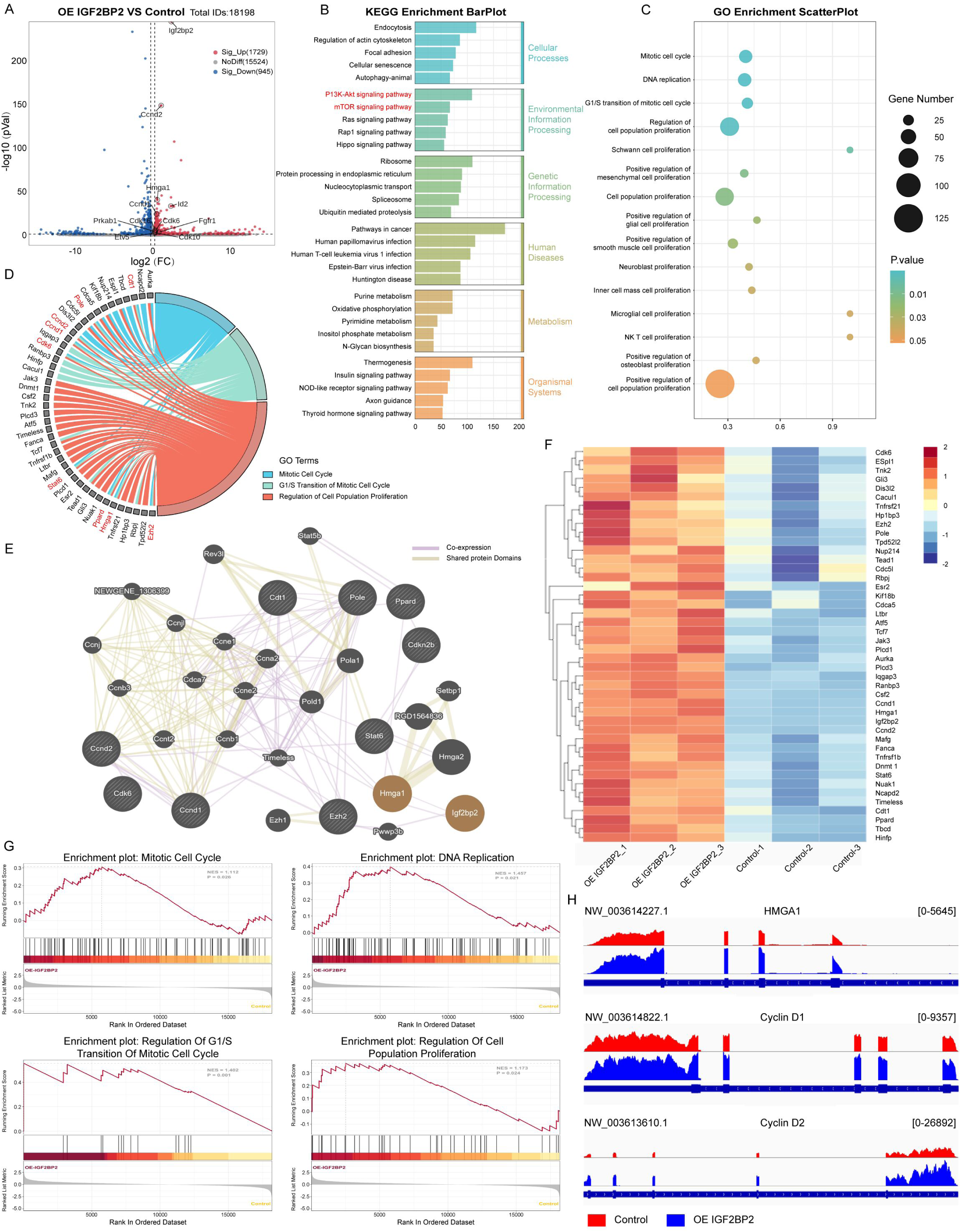
Gene expression changes, marked by activation of the HMGA1 gene and other proliferation-related genes, result from overexpression of the IGF2BP2 gene. (A) Volcano plot showing differentially expressed genes (DEGs) between OE IGF2BP2 and control cells. Highlighted points represent genes associated with proliferation, function, epigenetic modification, and the mitotic cell cycle. (B) Kyoto Encyclopedia of Genes and Genomes (KEGG) analysis of all DEGs in OE IGF2BP2 versus control CHO cells. (C) Gene Ontology (GO) analysis of proliferation-related genes in OE IGF2BP2 cells compared to control CHO cells. (D) GO enrichment circle diagrams display representative pathways and associated genes. Outer left track: list of representative genes. Outer right track: corresponding GO terms, each distinguished by a specific color. Three terms are shown, corresponding to up-regulated GO categories from RNA-seq analysis of OE IGF2BP2 versus control CHO cells. (E) Protein–protein interaction (PPI) network of differentially expressed genes from GeneMANIA analysis of genes in panel D. (F) Heatmap showing representative proliferation-related DEGs and their fold upregulation compared to the control. (G) Gene Set Enrichment Analysis (GSEA) revealed significant upregulation of the mitotic cell cycle, DNA replication, regulation of G1/S transition of the mitotic cell cycle, and regulation of cell population proliferation in OE IGF2BP2 cells compared to controls. P-values were calculated using Fisher’s exact test. (H) IGV tracks show RNA-seq signals for HMGA1, Cyclin D1, and Cyclin D2 transcripts.

### 4. Effects of HMGA1-KD on recombinant protein expression in stably transfected CHO cells

To investigate the role of HMGA1 in CHO cells, we constructed three distinct shRNA plasmids (shHMGA1-1, -2, and -3) and transfected these into CHO cells. After selection with G418, we established stable HMGA1-knockdown cell lines. Quantitative PCR (qPCR) analysis showed significant reductions in HMGA1 mRNA levels: 39.33% ± 0.04 (shHMGA1-1), 22% ± 0.02 (shHMGA1-2), and 40.33% ± 0.04 (shHMGA1-3) compared to the control (Figure 4A). Western blot analysis further confirmed decreases in HMGA1 protein expression by 22.67% ± 0.04, 20.66% ± 0.03, and 37.67% ± 0.03, respectively (Figure 4B). CCK-8 assays and colony formation assays demonstrated that all knockdown cell lines inhibited CHO cell proliferation (Figure 4C-E). Immunofluorescence similarly indicated higher knockdown efficiency for shHMGA1-2 (Figure 4F,G) which was selected for further study (termed HMGA1 KD). Three recombinant protein expression vectors (rADM, rVN, and rDONA) were transferred into the experimental and control groups for suspension culture, respectively. Results showed HMGA1 knockdown significantly inhibited recombinant proteins production, reducing titers (rADM 50.27%±0.03, rVN 70.89%±0.08, rDONA 38.17%±0.07) and specific productivity (rADM 53.7%±0.02, rVN 76.43%±0.01, rDONA 40.16%±0.02) compared to controls. Concurrently, viable cell density was significantly reduced (Figure 4H-M).

**Figure 4.**
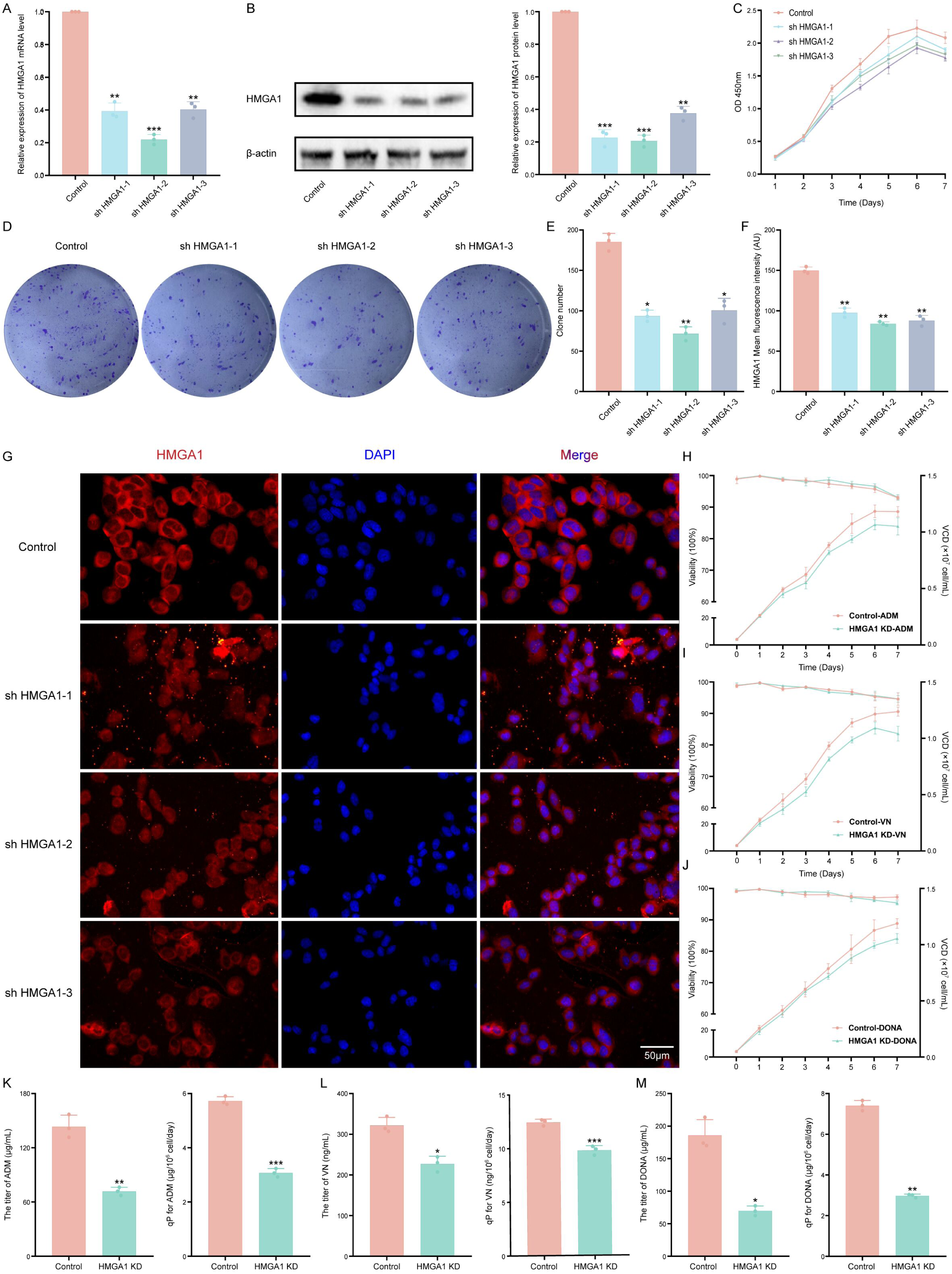
Effects of HMGA1 knockdown (KD) on recombinant protein expression in stably transfected CHO cells. (A) HMGA1 mRNA expression was measured using qPCR, while its protein levels were determined by Western blot analysis (B). (C) Cell proliferation was assessed via the CCK-8 assay. (D, E) Clonogenic potential was evaluated using a colony formation assay. (F) Immunofluorescence staining was performed on CHO cells with an anti-HMGA1 antibody (red); nuclei were counterstained with DAPI (blue). The average fluorescence intensity (in arbitrary units, AU) was also calculated (G). Viability and viable cell density (VCD) of CHO cells were examined following exposure to rADM (H), rVN (I), or rDONA (J) across both groups. The titer and quantitative production (qp) of rADM (K), rVN (L), and rDONA (M) were assessed by ELISA in the two experimental groups. (*p < 0.05; **p < 0.01; ***p < 0.001)

### 5. m6A modification of HMGA1 mRNA maintains its stability in an IGF2BP2-dependent manner

Previous studies have shown that IGF2BP2 regulates HMGA1 mRNA expression ^16^. To determine if IGF2BP2 influences HMGA1 post-transcriptional modification in CHO cells, we performed MeRIP-qPCR. This analysis revealed a significant increase in m6A-modified HMGA1 mRNA upon IGF2BP2 overexpression (Figure 5A). Protein stability assays further showed that IGF2BP2 overexpression does not alter HMGA1 protein stability (Figure 5B). However, RNA stability assays demonstrated that HMGA1 mRNA stability is enhanced in IGF2BP2-overexpressing CHO cells (Figure 5C).

**Figure 5.**
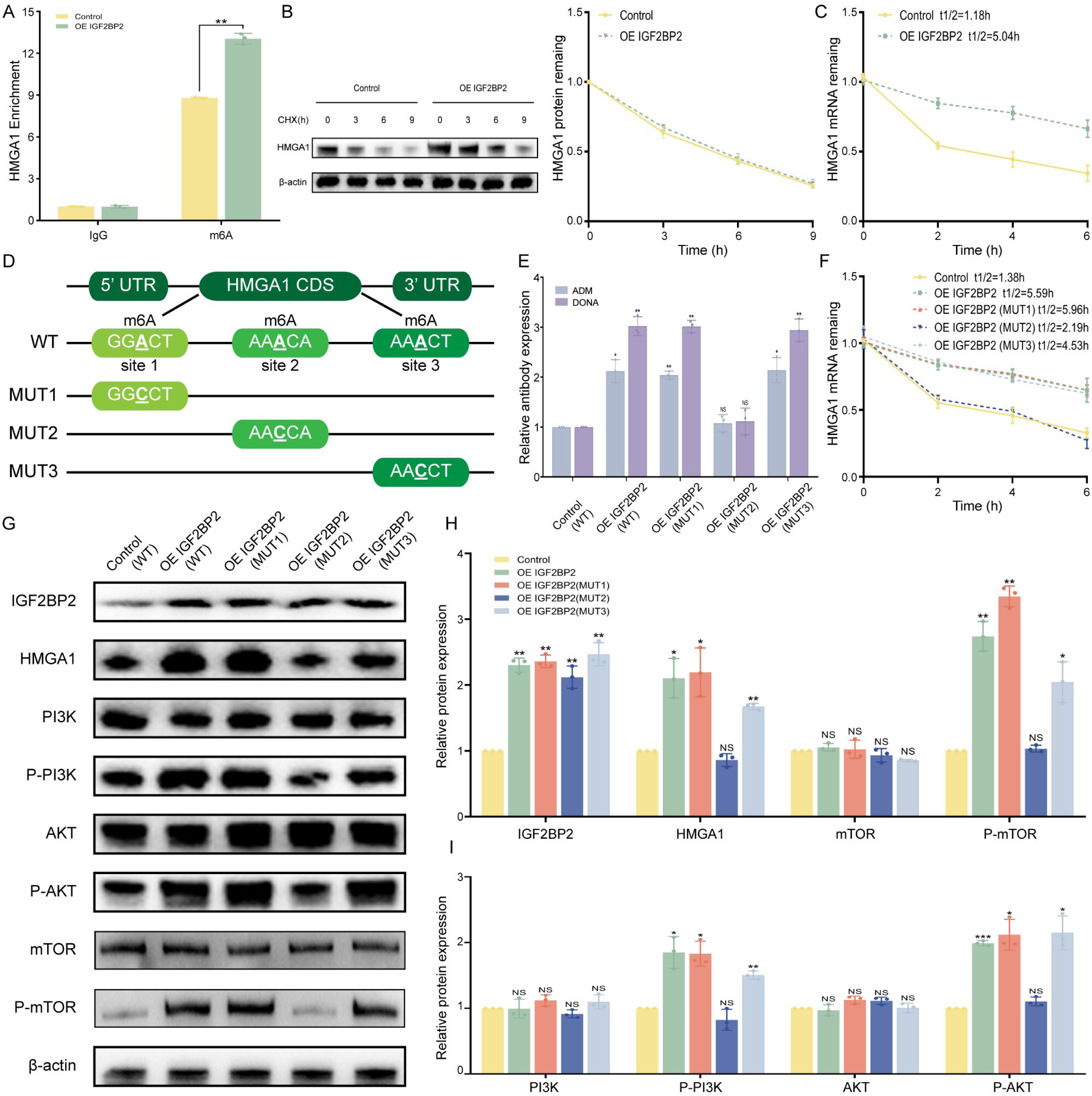
IGF2BP2 enhances HMGA1 m6A modification and mRNA expression. (A) MeRIP-qPCR analysis confirmed that the levels of m6A peaks on HMGA1 mRNA are modulated by IGF2BP2 overexpression. (B) A protein stability assay was performed on IGF2BP2-overexpressing (OE) and control cells treated with cycloheximide (CHX) at the indicated time points. (C) RNA stability was assessed in IGF2BP2-overexpressing (OE) and control cells after treatment with actinomycin D for various time periods, with quantitative polymerase chain reaction (qPCR) used to evaluate HMGA1 mRNA expression levels. (D) This study examined potential m6A modification sites in the HMGA1 coding sequence (CDS), and synonymous mutations were introduced into the HMGA1 CDS. (E) The relative expression levels of rADM and rDONA were measured by ELISA in different mutant vectors. (F) This study aimed to determine the effects of different mutant vectors on HMGA1 half-life stability. (G) Western blot analysis demonstrated that mutant 2 reversed the downstream pathway activation induced by IGF2BP2 overexpression. (p < 0.05; **p < 0.01; ***p < 0.001)

Since IGF2BP2 regulates RNA metabolism by stabilizing its target mRNAs, we sought to identify the specific m6A sites on HMGA1 recognized by IGF2BP2. Based on a previous report (Wang, et al, 2025), we predicted m6A sites and constructed a wild-type (WT) plasmid containing the full coding sequence (CDS), and three mutant plasmids (Mut1, Mut2, Mut3). Each mutant plasmid contained an A-to-C point mutation designed to abolish m6A methylation at a specific site (Figure 5D). Given the known positive correlation between HMGA1 expression and recombinant monoclonal antibody production, we introduced these mutants into the OE IGF2BP2 group and measured antibody expression by ELISA. The results showed that mutation at site 2 (Mut2) significantly reduced antibody expression. This result was supported by RNA stability assays, which showed decreased HMGA1 mRNA stability for Mut2 (Figure 5E, F). Western blot analysis of relevant pathways after the mutation showed that Mut2 also downregulated the PI3K/AKT signaling pathway, which was initially activated by IGF2BP2 overexpression (Figure 5G, H). Together, these findings indicate that HMGA1 expression is regulated by IGF2BP2-mediated m6A modification.

### 6. HMGA1-mediated regulation of the PI3K/AKT/mTOR pathway in CHO cell proliferation

HMGA1 promotes cell proliferation through PI3K/AKT-dependent mechanisms and activates multiple molecular signaling pathways linked to tumor progression (Bian et al, 2022; Liau et al, 2008). To test this, we performed confocal microscopy on the Control group, the OE IGF2BP2 group, and the OE IGF2BP2 group treated with the PI3K/AKT pathway inhibitor T60564. The results indicated that IGF2BP2 overexpression activates the PI3K pathway, an effect that is inhibited by T60564 (Figure 6A). Further experiments showed that adding the PI3K/AKT agonist T13376 to the Control group significantly increased protein titers (reaching 181.22%±0.05 and 140.05%±0.04 of the control rADM titer at 20 μM), specific productivity (qP) (reaching 154.95%±0.04 and 126.15%±0.02 of the control rADM qP at 20 μM), as well as live cell density and viability (Figure 6B-E). We then treated the Control and OE IGF2BP2 groups with small molecule agonists and inhibitors targeting both the PI3K/AKT pathway and the mTOR pathway (agonist T14033 and inhibitor T1537). Western blot analysis of downstream pathway components confirmed that IGF2BP2 regulates the PI3K/AKT/mTOR pathway via HMGA1 (Figure 6F).

**Figure 6.**
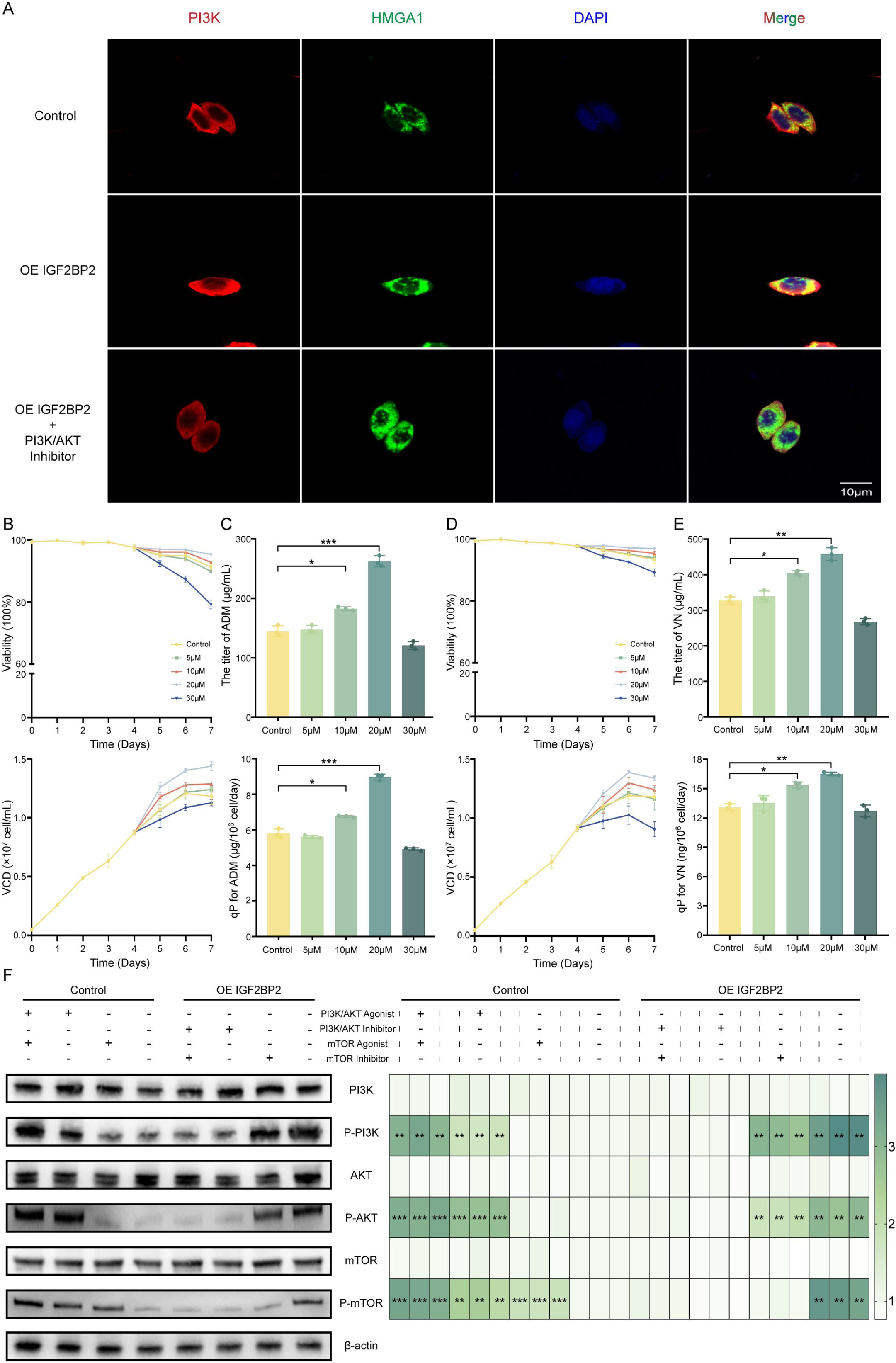
HMGA1-mediated regulation of the PI3K/AKT/mTOR pathway in CHO cell proliferation. (A) Laser confocal microscopy reveals a significant positive correlation between IGF2BP2 and HMGA1 expression and activation of the PI3K signaling pathway, with HMGA1 serving as a downstream effector of IGF2BP2. Scale bar: 10 µm. Pharmacological activation of the PI3K/AKT signaling pathway enhances recombinant protein production in CHO cells. Cell viability and VCD during the 7-day culture period with the PI3K/AKT agonist (T13376) at concentrations of 5, 10, 20, and 30 µM are shown in panels (B) and (D), respectively. (C, E) ELISA quantification of rADM/rVN expression levels and total antibody titers. (*p < 0.05; **p < 0.01; ***p < 0.001). (F) Western blot analysis of IGF2BP2 expression in the OE IGF2BP2 group versus the control group, following treatment with small-molecule activators and inhibitors of the PI3K/AKT and mTOR pathways, demonstrating IGF2BP2’s role in modulating downstream signaling. (*p < 0.05; **p < 0.01; ***p < 0.001)

### 7. IGF2BP2 governs metabolic reprogramming in CHO cells by orchestrating antioxidant defense and mitochondrial homeostasis

Following the initial elucidation of the IGF2BP2/HMGA1 axis and its role in downstream proliferative signaling, we transfected enhanced green fluorescent protein (eGFP) into Control, OE IGF2BP2, and OE IGF2BP2+HMGA1 KD groups. Immunofluorescence analysis revealed the mean fluorescence intensity of IGF2BP2, HMGA1, and eGFP (Figure 7A, B). Interestingly, IGF2BP2-mediated HMGA1-induced host cell proliferation was not the sole driver of recombinant protein expression. ELISA further demonstrated recombinant antibody levels across cell line groups (Figure 7C), suggesting that IGF2BP2 may additionally regulate recombinant protein expression through an alternative pathway.

**Figure 7.**
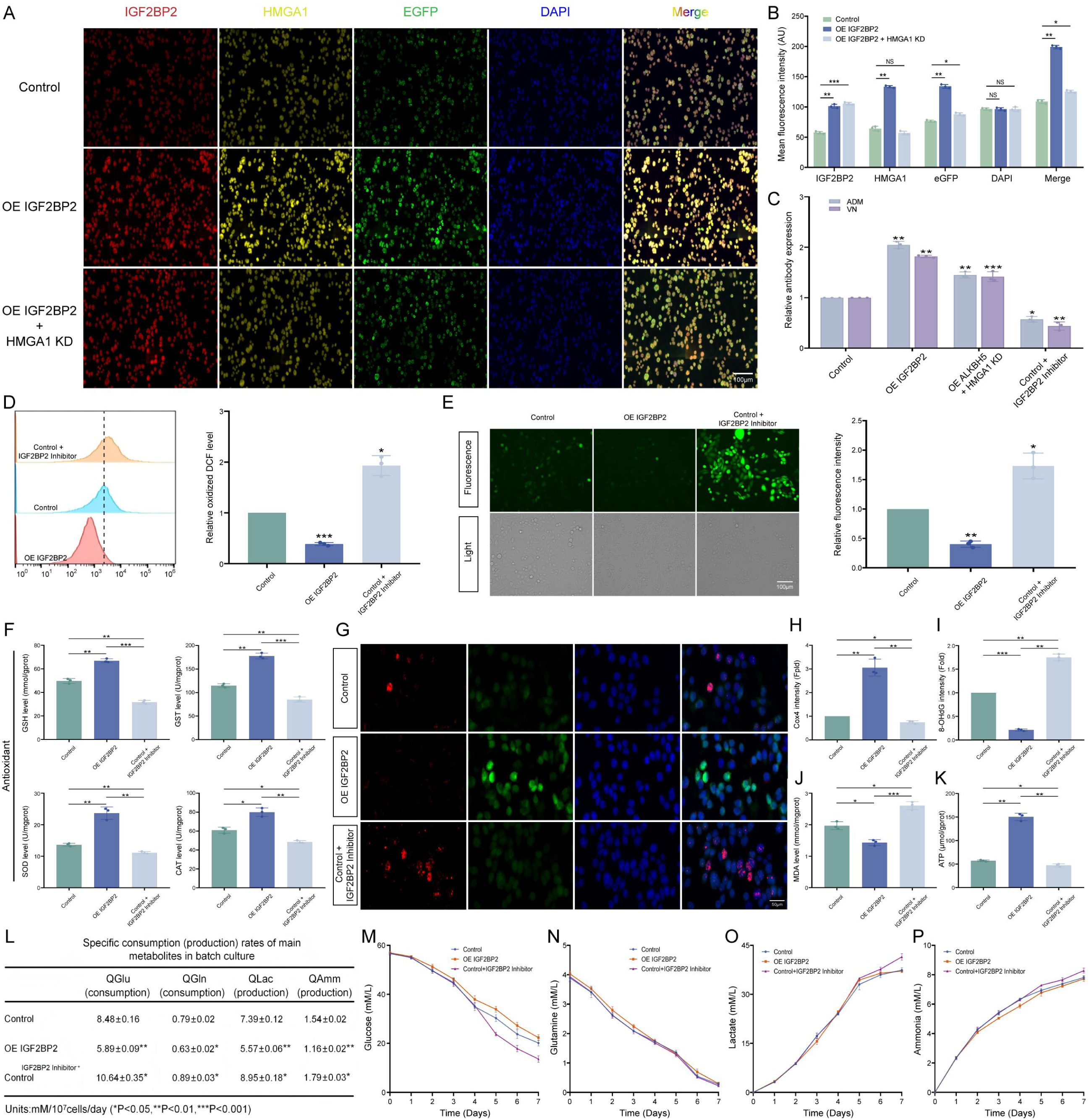
The role of IGF2BP2 in antioxidant defense and mitochondrial protection in CHO cells and its effects on metabolic characteristics. (A) A multicolor immunofluorescence assay demonstrated a significant positive correlation among IGF2BP2, HMGA1, and eGFP in the Control, OE IGF2BP2, and Control+IGF2BP2 Inhibitor groups (with IGF2BP2 labeled in red, HMGA1 in yellow, eGFP in green, and DAPI in blue). (B) Quantitative analysis of the average fluorescence intensity (in arbitrary units, AU) was also performed. (C) ELISA was used to measure the relative expression levels of rADM and rVN across various groups. (D) Reactive oxygen species (ROS) levels were assessed using the fluorescent probe DCFH-DA in the Control, OE IGF2BP2, and Control+IGF2BP2 Inhibitor groups, followed by quantitative analysis. (E) ROS levels were visualized by fluorescence microscopy (images are representative of three independent experiments) and quantified for fluorescence intensity using Image J software. (F) Antioxidant markers were measured, including glutathione (GSH) content, glutathione S-transferase (GST) activity, superoxide dismutase (SOD) activity, and catalase (CAT) activity. (G-I) Immunofluorescence analysis of COX4 and 8-OHdG levels. (J) Malondialdehyde (MDA) content was measured. (K) ATP content was analyzed. (L) Specific consumption/production rates (QI = Δ I(t2-t1)/ Δ VCD) were determined. (M-P) Temporal profiles of glucose, lactate, glutamine, and ammonia concentrations are shown. (*p < 0.05; **p < 0.01; ***p < 0.001)

Oxidative stress, mitochondrial function, and metabolic pathways are key factors that suppress CHO cell proliferation and function in long-term culture (Gao, et al, 2025; Zhang et al, 2025) and are closely associated with IGF2BP family regulation (Zeng, et al, 2025; Gao et al, 2024; Song et al, 2024). To determine whether IGF2BP2 influences host cell reactive oxygen species (ROS), we detected ROS using the DCFH-DA probe in Control, OE IGF2BP2, and Control+IGF2BP2 Inhibitor groups. Both flow cytometry and fluorescence microscopy indicated a significant negative correlation between IGF2BP2 expression and ROS levels (Figure 7D, E). Consistent with this, IGF2BP2 overexpression increased activities of key antioxidant enzymes—glutathione (GSH), glutathione S-transferase (GST), superoxide dismutase (SOD), and catalase (CAT)–by 1.35 ± 0.03, 1.55 ± 0.03, 1.73 ± 0.07, and 1.31 ± 0.09-fold relative to the control, respectively (Figure 7F). Moreover, elevated cytochrome c oxidase subunit IV (COX4) expression, alongside decreased oxidative damage markers 8-OHdG and malondialdehyde (MDA), indicated enhanced electron transport chain activity and reduced oxidative damage in IGF2BP2-overexpressing cells (Figure 7G–J). This was further supported by a significant increase in intracellular ATP levels upon IGF2BP2 overexpression (Figure 7K), suggesting improved mitochondrial energy production to support recombinant protein expression (Zhang et al, 2020).

Metabolite profiling during suspension culture revealed that the OE IGF2BP2 group had lower specific consumption rates of glucose and glutamine, as well as lower specific production rates of lactate and ammonia, compared with the control (Figure 7L–P). To further validate IGF2BP2’s metabolic role, an IGF2BP2 inhibitor was added during the logarithmic growth phase (Control+IGF2BP2 Inhibitor). Notably, the OE IGF2BP2 group exhibited higher ATP production and recombinant protein expression, indicating improved energy conversion and utilization efficiency–a promising trait for long-term cultivation. Reduced lactate and ammonia accumulation in this group aligns with reports that such metabolites impair recombinant antibody production by disrupting lactate metabolism and mitochondrial function (Badsha et al, 2016; Ishii et al, 2015), suggesting that enhanced recombinant protein expression may result from an activated TCA cycle and attenuated ammonia toxicity.

## Discussion

To improve the production efficiency of recombinant proteins in mammalian expression systems and address their instability in expression, numerous studies have investigated the role of epigenetic regulatory mechanisms (Zhang, et al, 2025; Li et al, 2022). Among these, DNA methylation and histone modifications have been shown to significantly increase the expression levels and stability of recombinant proteins, especially antibodies, in Chinese hamster ovary (CHO) cells (Yang et al, 2025). Additionally, epigenetic approaches have been used to enhance anti-apoptotic traits and metabolic pathways in cells, thereby supporting protein synthesis (Zhang, et al, 2025; Zhang et al, 2021). Researchers have also performed screening studies to explore the potential function of RNA modifications, such as m6A, in regulating gene expression. In this study, we revealed a marked difference in IGF2BP2 expression between CHO cell lines with high and low recombinant monoclonal antibody production. An IGF2BP2 overexpression model provided strong evidence that this factor promotes CHO cell proliferation. Following this, combined phenotypic analysis and transcriptome sequencing further identified HMGA1 as a key downstream target. Mechanistically, IGF2BP2 appears to enhance HMGA1 mRNA stability by recognizing and binding to specific m6A modification sites, which activates proliferation-related signaling pathways and ultimately improves CHO cell growth and recombinant antibody yield. Engineering strategies aimed at stimulating cell proliferation have proven effective in raising recombinant protein expression levels, primarily by increasing the overall number of productive cells and extending the duration of high-rate protein production (Zhang, et al, 2025).

High Mobility Group A (HMGA) proteins are a class of non-histone chromatin structural proteins that belong to the High Mobility Group (HMG) protein subfamily. They primarily regulate chromatin conformation and gene transcription by binding to AT-rich DNA sequences via their AT-hook domains, and play crucial roles in embryonic development, cell differentiation, proliferation, and diseases such as cancer (Singh et al, 2025; Nué-Martinez et al, 2024; Battista et al, 2024). Among these, HMGA1, as a core member, is overexpressed in various cancer cells, including colorectal cancer, ovarian cancer, and hematological malignancies. It promotes cell proliferation, invasion, and metastasis by regulating cell cycle-related genes, such as Cyclin D1 (E et al, 2020; Ma et al, 2021; Zhao et al, 2024). Similarly, in CHO cell bioprocess development, modulating cell proliferation has been proven to be a reliable strategy for enhancing recombinant protein productivity (Roshanmehr et al, 2024; Liu et al, 2022; Torres et al, 2023). Consequently, process optimization guided by proliferation enhancement represents a well-established and mechanistically clear pathway to intensify and stably increase product titers in CHO cell cultures. Research indicates that IGF2BP2 directly binds to and stabilizes HMGA1 mRNA in an m6A-dependent manner by forming a LINC00460/DHX9/IGF2BP2 ternary complex, thereby upregulating HMGA1 protein expression and driving proliferation, migration, and metastasis in colorectal cancer cells (Hou, et al, 2021). Simultaneously, evidence suggests a potential HMGA2-IGF2BP2-HMGA1 positive feedback regulatory loop, wherein IGF2BP2 transcription activation further amplifies this pro-proliferative mechanism (Sun et al, 2022). This study similarly demonstrates that IGF2BP2 is conserved in CHO cells, where it enhances cellular proliferation by activating downstream HMGA1 m6A hyperactivation sites. It is worth noting that enhanced cell proliferation is not the only mechanism through which IGF2BP2 increases recombinant antibody expression in CHO host cells. Investigations into its effects on overall cellular metabolic activity reveal that IGF2BP2 reduces oxidative stress, improves mitochondrial function, enhances the utilization of glucose and glutamine, and partially mitigates the accumulation of lactate and ammonia. Research in neural stem cells demonstrates that IGF2BP2 overexpression significantly boosts mitochondrial membrane potential, ATP production, and cellular oxygen consumption rate (Zeng, et al, 2025). Similarly, studies indicate that IGF2BP3 stabilizes NFAT5 via an m6A-dependent mechanism, indirectly optimizing mitochondrial function and increasing ATP output (He, et al, 2025). These findings are corroborated by the experimental results of this study: elevated ATP levels, increased expression of electron transport chain components (e.g., COX4), and a marked decrease in the oxidative damage marker 8-OHdG. The enhancement of these metabolic traits suggests that genetic engineering of host cell lines or improvements in culture conditions can directly influence cell growth dynamics, thereby further enhancing production efficiency. Specifically, optimizing subculture protocols can produce significant outcomes. For instance, using an optimized Super7 subculture strategy to mimic production phase conditions enables cells to sustain superior net growth rates and oxidative metabolism during extended culture, ultimately increasing monoclonal antibody production in unstable cell lines by 80% to 160% (Moran et al, 2024). Furthermore, metabolic modeling studies show that cell growth status is tightly linked to the core metabolic network. Optimizing proliferation conditions facilitates the coordination of key metabolic fluxes (such as amino acid cycling reactions), thereby promoting efficient protein synthesis (Jiménez Del Val et al, 2023).

In conclusion, this study demonstrates that targeted genetic editing of CHO cell lines can promote host cell proliferation and metabolic remodeling, concurrently enhancing the specific growth rate and specific yield of recombinant proteins, ultimately resulting in a significant increase in production efficiency. This further confirms that interventions in CHO cell growth dynamics can be effectively translated into gains in final production efficiency, supporting the proliferation-production coupling effect at the cellular physiological level. Future work should involve the long-term cultivation of engineered cell lines in bioreactors to comprehensively assess their growth, production efficiency, stability, metabolic profiles, and the critical quality attributes of recombinant proteins. Similarly, validation across multiple cell lines and yield confirmation are of equal importance. Despite numerous research avenues remaining for future exploration, this study has, for the first time, unveiled the critical role of the IGF2BP2/HMGA1/mTOR axis in enhancing host cell proliferation. Additionally, it has elucidated how IGF2BP2 influences cell metabolic reprogramming by ameliorating oxidative stress and mitochondrial function.

## Conclusion

In summary, this study reveals a novel epitranscriptomic regulatory mechanism in Chinese Hamster Ovary (CHO) cells: the RNA-binding protein IGF2BP2 specifically recognizes and stabilizes m6A modification sites on HMGA1 mRNA, thereby activating the downstream PI3K/AKT/mTOR signaling pathway. IGF2BP2 not only significantly promotes cell proliferation and the yield of recombinant antibody proteins but also comprehensively reshapes cellular redox homeostasis and metabolic capacity by enhancing the cell’s antioxidant defense and promoting mitochondrial ATP production and utilization. These findings elucidate a new paradigm wherein an m6A methylation reader co-regulates mitochondrial function and cell proliferation to drive recombinant protein production, providing an innovative epigenetic strategy for constructing high-yielding CHO cell lines. Targeting IGF2BP2 or the m6A epitranscriptome holds promise as a novel approach to optimize bioprocesses, and its translational potential awaits further validation in future industrial-scale production systems.

## Materials and methods

### 1. Cell culture and reagents

CHO-S cells (Catalog #A11557-01; Life Technologies, Carlsbad, CA, USA), together with recombinant CHO cell clones displaying either high or low productivity of adalimumab (ADM), recombinant human vitronectin (rVN), recombinant adalimumab (rADM), and recombinant donanemab (rDONA), were sourced from the International Joint Research Laboratory for Recombinant Pharmaceutical Protein Expression System (Xinxiang, China). The recombinant protein-producing clones, categorized as ADMH (high productivity) and ADML (low productivity) for ADM variants, were developed using limited dilution cloning technology (Gao et al, 2025; Yang et al, 2022; Lu et al, 2025). Suspension cultures were set up at an initial density of 5.0 × 10⁵ cells/mL in 30 mL of AF-CD medium (ProteinEasy, Xinxiang, China) placed in 125-mL conical flasks (Biosharp, Anhui, China). The cultures were continuously shaken at 110 rpm and maintained under standard incubation parameters (37°C, 85% humidity, 5% CO₂). Detailed information on the small-molecule additives and related assay kits such as ATP and MDA used in this experiment is provided in Table 1.

**Table 1:** Specific information on small-molecule additives and related detection kits

| Name | Manufacturer |
| --- | --- |
| IGF2BP2 inhibitor | T67930L; TargetMol, China |
| PI3K/AKT inhibitor | T60564; TargetMol, China |
| mTOR inhibitor | T1537; TargetMol, China |
| PI3K/AKT activator | T13376; TargetMol, China |
| mTOR activator | T14033; TargetMol, China |
| ATP assay kit | A095-2-1, Nanjing Jiancheng Bioengineering Institute, China |
| MDA assay kit | A003-4-1, Nanjing Jiancheng Bioengineering Institute, China |
| GSH assay kit | A006-1-1, Nanjing Jiancheng Bioengineering Institute, China |
| GST assay kit | A004-1-1, Nanjing Jiancheng Bioengineering Institute, China |
| SOD assay kit | A001-3-1, Nanjing Jiancheng Bioengineering Institute, China |
| CAT assay kit | A007-1-1, Nanjing Jiancheng Bioengineering Institute, China |

### 2. Vector construction and transfection

The piggyBac vector (Haixing Biotechnology Co., Ltd.) served as the backbone for genetic modifications. Synthesized nucleic acid fragments were inserted downstream of the CAG promoter via restriction enzyme digestion and ligation to generate the IGF2BP2 overexpression vector (GenBank ID: 100755568). Concurrently, HMGA1 knockdown vectors (shHMGA1-1 to -3) were constructed by Starfish Biotechnology Co., Ltd., with the following shRNA sequences: shHMGA1-1: 5′-AGTCAGAAGGAGCCCAGTGAA-3′, shHMGA1-2: 5′-CAGACCCAAGAAACTGGAGAA-3′, shHMGA1-3: 5′-GGACGGGACTGAGAAGCGAGG-3′. CHO cells were transfected at 75% confluence using Lipofectamine 2000 in Opti-MEM medium according to the manufacturer’s protocol. After 8 hours, the medium was replaced with DMEM/F12 containing 10% FBS. Stable transfected pools were selected with antibiotics and harvested after two weeks of continuous culture.

### 3. RT-qPCR analysis of mRNA expression levels

Quantitative reverse transcription PCR (qPCR) was performed to measure IGF2BP2 mRNA expression in CHO cells across experimental groups: control, overexpression. Gene-specific primers for IGF2BPs and related targets were designed using NCBI database references and vector sequences (Table 2). Briefly, cells were lysed directly with TRIzol reagent (Biosharp, Anhui, China) containing RNase inhibitors, followed by total RNA extraction and reverse transcription into cDNA. Amplification reactions were carried out in a 20 μL system containing SYBR Green premix (Vazyme, Nanjing, China) with ROX reference dye, using a three-step cycling protocol (denaturation, annealing, extension). The 2^−ΔΔCT^ method was employed for data analysis, with target gene Ct values normalized to the geometric mean of housekeeping genes.

**Table 2.**
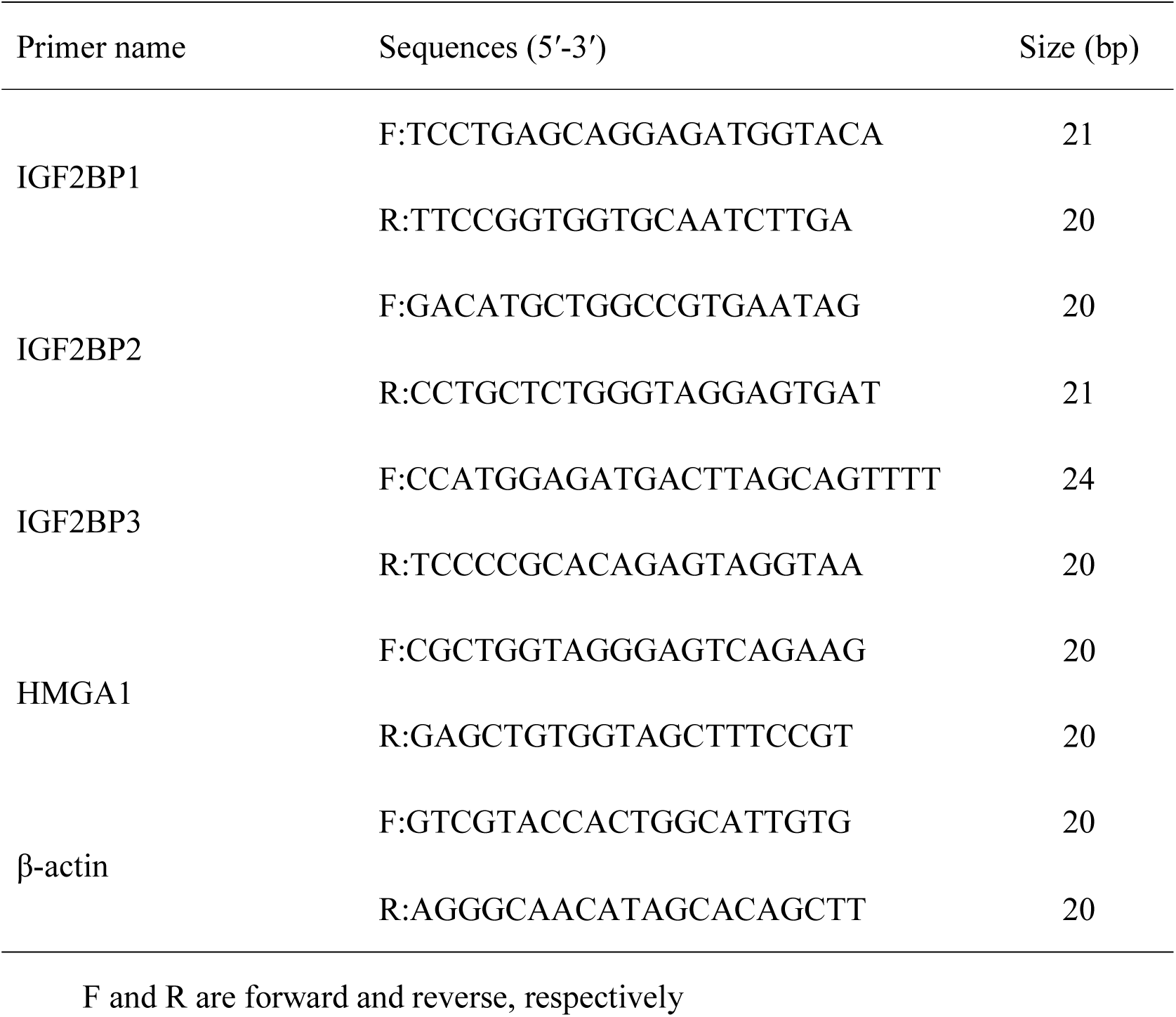
Sequences of the primers for quantitative real-time PCR

### 4. Western blotting and protein extraction procedures

Protein levels of IGF2BP2 and HMGA1 were detected in different cell lines using Western blotting. For intracellular protein analysis, total proteins were extracted using RIPA lysis buffer with protease inhibitors. Protein concentration was determined by BCA assay and normalized. Proteins were separated by 4%–12% SDS-PAGE, transferred to PVDF (Sigma Aldrich, St. Louis, MO, USA) membranes, blocked, and incubated with primary antibodies. β-actin was used as a loading control. After TBST washes, membranes were incubated with HRP-conjugated secondary antibodies and detected using ECL chemiluminescence. Suspended cell supernatants were processed similarly, with anti-human IgG secondary antibody (1:10,000 dilution) (E030170-01, EARTHox, USA). Antibody details are listed in Table 3.

**Table 3.** Antibodies used in this study.

| Antibodies | Antibodies | Concentration |
| --- | --- | --- |
| Anti-IGF2BP2 | 11601-1-AP, Proteintech | 1/2000(WB), 1:400 (IF) |
| Anti-β-actin | 66009-1-Ig, Proteintech | 1/1000(WB) |
| Anti-HMGA1 | Ab315350, Abcame | 1/1000(WB), 1:400 (IF) |
| Anti-mTOR | AF6308, Affinity | 1/1000(WB) |
| Anti-P-mTOR | AF3309, Affinity | 1/1000(WB) |
| Anti-PI3K | A18355, Abclonal | 1/1000(WB), 1:100 (IF) |
| Anti-P-PI3K | AP1463, Abclonal | 1/1000(WB) |
| Anti-AKT | A18675, Abclonal | 1/1000(WB) |
| Anti-P-AKT | AP1453, Abclonal | 1/1000(WB) |
| Anti-Cox4 | A11631, Abclonal | 1:150 (IF) |
| Anti-8-OHdG | sc-66036, Santa Cruz<br>Biotechnology | 1:200 (IF) |

### 5. Immunostaining

Pre-cleaned coverslips were placed in a 6-well plate, and each was seeded with 1 mL of cell suspension (1-2 × 10⁴ cells). After 4 hours of adhesion, 1 mL of complete medium was added per well, followed by overnight incubation at 37°C with 5% CO₂. Cells were fixed using 4% PFA for 10–15 minutes at room temperature. Primary antibody incubation followed overnight at 4°C according to Table 3. Secondary antibody (Invitrogen, Carlsbad, CA, USA) (Alexa Fluor 488-conjugated) was applied for 1 hour in the dark. Nuclei were stained with DAPI for 5 minutes. Fluorescence imaging was performed using appropriate filters.

### 6. Cellular proliferation, colony formation assays

#### CCK-8 Kinetic Analysis

Pretreated cells were seeded in 96-well plates. At indicated time points (24–144 h), 10 μL of CCK-8 reagent (Topscience, China) was added per well. After 1 h of incubation at 37°C in the dark, absorbance was measured at 450 nm using a microplate reader (Molecular Devices, USA). Controls (medium-only and reagent-only wells) were included for normalization.

#### Clonogenic Formation Assay

Genetically modified cells were plated at low density (500 cells/well) in 6-well plates and cultured for 10 days. Colonies were fixed with methanol, stained with 0.5% crystal violet (Beyotime, China), and counted. Plating efficiency was calculated as (number of colonies / cells seeded) × 100%.

### 7. Analysis of Apoptosis and Cell Cycle

Apoptosis Analysis: Cells in calcium-containing binding buffer were stained with Annexin V-APC (1:20) and 5 μg/mL DAPI, incubated at 37°C in dark (5/15/30 min intervals), and immediately chilled (4°C) to halt staining. Flow cytometry used 488 nm (DAPI) and 640 nm (APC) lasers, with FSC/SSC gating to exclude debris.

Cell Cycle Analysis: Log-phase cells were fixed in chilled 70% ethanol (4°C, 2 h), stained with PI/RNase A (100 μg/mL, 37°C, 40 min in dark), and analyzed by flow cytometry (405 nm excitation, low flow rate <300 events/sec). Pulse width gating excluded aggregates. Data were processed using ModFit LT (FlowJo v10.8) for phase quantification.

### 8. Enzyme-Linked Immunosorbent Assay (ELISA)

The collected supernatant from the suspension cell cultures was centrifuged to prepare the samples. We detected ADM and DONA antibody levels, as well as VN protein yield, using the Human IgG1 ELISA Kit (Ruixinbio, Quanzhou, China) and the Vitronectin (VN) ELISA Kit (Ruixinbio, Quanzhou, China). Specific yield was calculated as 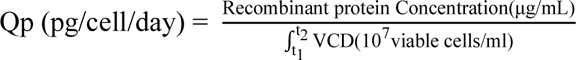

### 9. RNA-Seq and analysis

For eukaryotic samples, mRNA was enriched via poly(A)+ selection and fragmented into approximately 300-bp segments. High-fidelity reverse transcriptase with Unique Molecular Identifiers (UMIs) was used for cDNA synthesis to minimize template loss and correct PCR bias. The resulting PCR products were aligned to the reference genome using HISAT2, followed by 150-bp paired-end sequencing on the Illumina NovaSeq 6000 platform. Differential gene expression analysis was performed with DESeq2, while Gene Ontology and KEGG pathway analyses were conducted using clusterProfile (Lu, et al, 2025).

### 10. Methylated RNA immunoprecipitation (MeRIP)-qPCR

MeRIP assays were performed with a commercial MeRIP m6A kit (Epigentek, USA). Total RNA was isolated from cultured cells using Trizol reagent, followed by DNase I treatment to eliminate genomic DNA contamination. Magnetic beads coated with protein A/G were pre-incubated with anti-m6A antibody (Epigentek, USA) or anti-IgG (as a negative control) at room temperature for 90 minutes. The RNA was then incubated with the antibody-bound beads at 4 °C to facilitate m6A-specific binding. Bead-bound complexes were subsequently isolated using a magnetic stand and washed thoroughly with washing buffer. Finally, bound RNA was eluted and subjected to reverse transcription and quantitative PCR following the procedures detailed in Section 3 of Materials and Methods.

### 11. RNA/Protein stability assays

RNA stability Assay: Following established methodologies, CHO cells were treated with 5 μg/mL actinomycin D (TargetMol, Shanghai) to induce transcriptional arrest. Total RNA was extracted at predefined time intervals and reverse-transcribed into cDNA. The RNA decay rate was subsequently determined via quantitative PCR (qPCR) and analyzed according to previously described computational models (Blumberg et al, 2021).

Protein stability assays: CHO cells seeded in 6-well plates were exposed to 100 μg/mL cycloheximide (TargetMol) to inhibit protein synthesis. Whole-cell lysates were collected at specified time points, and the degradation kinetics of HMGA1 were evaluated by Western blotting. Half-life calculations were derived from band intensity quantitation.

### 12. Statistical Analysis

All data are presented as mean ± standard deviation. Statistical analyses were performed using GraphPad Prism software (version 8.01, GraphPad Software, USA). Differences between groups were assessed by one-way ANOVA or unpaired two-tailed Student’s t-test, as appropriate. A p-value of less than 0.05 was considered statistically significant, with specific thresholds denoted as follows: *p < 0.05, **p < 0.01, and ***p < 0.001.

## Data availability statement

All relevant data are available from the authors. Data could be accessed by SRA series number PRJNA1397464.

## CREDIT Author Statement

Zhao-Ming Cui: Conceptualization, Methodology, Writing - Original Draft, Project administration. Jiang-Tao Lu: Conceptualization, Methodology, Writing - Review & Editing. Yuan-Ye Xu: Methodology, Validation. Ying-Ying Feng: Software, Data Curation, Visualization. Yang Guo: Investigation. Yan-Ping Gao: Investigation. Wen Wang: Formal analysis. Le-Le Qiu: Data Curation. Xiao-Yin Wang: Visualization, Funding acquisition. Zi-Chun Hua: Methodology, Writing - Review & Editing. Tian-Yun Wang: Funding acquisition. Yan-Long Jia: Resources, Funding acquisition.

## Ethics approval statement

This article does not contain any studies with human or animal subjects

